# Impact of the Excitatory-Inhibitory Neurons Ratio on Scale-Free Dynamics in a Leaky Integrate-and-Fire Model

**DOI:** 10.1101/2023.11.28.569071

**Authors:** Mohammad Dehghani-Habibabadi, Nahid Safari, Farhad Shahbazi, Marzieh Zare

## Abstract

The relationship between ratios of excitatory to inhibitory neurons and the brain’s dynamic range of cortical activity is crucial. However, its full understanding within the context of cortical scale-free dynamics remains an ongoing investigation. To provide insightful observations that can improve the current understanding of this impact, and based on studies indicating that a fully excitatory neural network can induce critical behavior under the influence of noise, it is essential to investigate the effects of varying inhibition within this network. Here, the impact of varying ratios on neural avalanches and phase transition diagrams, considering a range of control parameters in a leaky integrate-and-fire model network, is examined. Our computational results show that the network exhibits critical, sub-critical, and super-critical behavior across different control parameters. In particular, a certain ratio leads to a significantly extended dynamic range compared to others and increases the probability of the system being in the critical regime. To address differences between various ratios, we utilized the Kuramoto order parameter and conducted a finite-size scaling analysis to determine the critical exponents associated with phase transitions. In order to characterize the criticality, we examined the distribution of neuronal avalanches at the critical point and the scaling behavior characterized by specific exponents.

## Introduction

The balance between excitatory and inhibitory neuronal activity, commonly referred to as the E/I balance, is a fundamental feature of neural network operation that has a significant impact on network dynamic stability, modulation of neural responses, and facilitating information processing^1–3^. The balance of these synaptic membrane currents is widely recognized as essential for the spontaneous activity of neurons and their responses to sensory stimuli^4–10^. On the other hand, disturbances of the cortical E/I balance in the brain emerged as a contributor to neurological diseases, such as epilepsy^11^, Parkinson’s disease (PD)^12^, Tourtette’s syndrome^13^, autism^14^, and schizophrenia^15–17^.

Although synaptic plasticity can balance a neuron’s input after a synaptic connection^18,19^, hyper-excitation could theoretically occur if the neural network has a significantly higher proportion of excitatory neurons than inhibitory neurons. On the other hand, an excess of inhibitory neurons can inhibit normal brain activity, resulting in reduced responsiveness or diminished functionality of neural circuits.^1,20,21^ Therefore, maintaining an optimal ratio of excitatory and inhibitory neurons is critical to ensuring a balanced interplay of excitation and inhibition in the brain. In addition, experimental studies have shown that approximately 20% of neurons are inhibitory GABAergic neurons in the cortex^22–28^.

The activities of inhibitory neurons have been identified as the source of various oscillatory rhythms^29^, and pharmacologically blocking them causes an epileptic seizure in cortical activity^30^. Moreover, it has been shown that excitation and inhibition are balanced in detail and globally^1,2,4,5,18,31^, providing a basis for neuronal oscillations and avalanches^32–34^. A neural avalanche refers to a sequential propagation of neuronal firing activity within a network in a correlated manner, initiated by the firing of a single neuron or a small group of neurons. The neural avalanche dynamic may critically depend on the E/I balance, as shown in experiments in cortical cultures, anesthetized rats, and awake monkeys^35–37^.

Neuronal oscillations and neural avalanches exhibiting global behavior embedded in neuronal activity patterns were observed *in vitro*^32,36,38^ and *in vivo*^39^. Numerous studies show that optimal operations including phase synchrony^36^, information storage^40^, communication, and information transition^32^, optimal communication^32,34,41–44^, transition capability^34^, computational power^42^, and dynamic range^34,45^ are found when the brain works near criticality. A critical point sets a boundary between an ordered and a less ordered state with different scaling behaviors^38^. At criticality, neural networks display power law distribution in a statistical collective behavior of the system, for instance, neural avalanche.

Studies show that avalanche size, and lifetime distributions are indicators of long-range correlation^32^. The widely varying profile of neural avalanche distribution in size is described by a single universal scaling exponent, *β* in size, *P*(*S*) ∼ *S*^*β*^, and a single universal exponent in duration, *τ*, *P*(*T*) ∼ *T*^*τ*^, where *S* and *T* denote the size, and the duration of avalanches, respectively. Power-law distributions in neuronal avalanches appeared in cortex slice cultures of rats in vitro^35,36^, rat cortical layer 2/3 at the beginning and end of the second week postnatal^46^, Local Field Potentials (LFPs) of anesthetized cats^47,48^, newborn rats^36^, and monkeys^39,48^.

Despite numerous experiments and modeling studies aimed at establishing the plausibility of the critical brain state and exploring the E/I ratio as a fundamental mechanism in brain dynamics, the impact of inhibition on both dynamics remains to be further investigated.

In this study, we address this problem by quantitatively analyzing the properties of a phase transition, including the order of the transition and its dynamic range, by varying the inhibition percentages systematically.

The order parameter is crucial in describing the transition between states in both critical equilibrium phenomena, e.g. Ising magnetic, which is a disordered-ordered transition^49^, and systems out of equilibrium phase, e.g. neural systems, which is a desynchronized-synchronized transition^50–52^ defined based on the Kuramoto order parameter^53–55^. In addition, the possibility of criticality based on the desynchronized-synchronized transition has been confirmed in magnetoencephalography (MEG) and intracranial stereoelectroencephalography (SEEG) during resting brain activity^56^.

In particular, we recently introduced the Kuramoto-based order parameter spike-phase coupling (*SPC*), which can detect the different levels of network-level synchrony in a purely excitatory leaky integrate-and-fire network^57^. Here, the *SPC* refers to the coupling between spike sequences^58,59^ and the phase of temporal fluctuations of a macroscopic observable of the network, namely the population average voltage (*PAV*)^60^. The study showed that an estimation of the strength of *SPC*, represented as ’*m*’, is equal to the average of the instantaneous phases of *PAV* for all spikes in the firing time series throughout the entire neuronal network^57^. The m-value is zero when synaptic efficacy or control parameter, *K*, is minimal and reaches one when it is substantial, resulting in complete synchronization of the network^57^.

In this study, we take into account the inhibition in the network and show a significant level of inhibition can prevent the network from being synchronized. In particular, when there is a dominance of excitation in a network with few or no inhibitions, the changes in *m* with respect to *K* are usually the same. This means that a significant value for *K* can result in full synchronization (*m* ≈ 1) in the network. In contrast, when inhibition is the dominant factor, complete synchronization is impossible, and *m* saturates at values below one, regardless of increasing *K*. As inhibition increases, *m* remains the same at a small value for all levels of control parameter.

Moreover, in line with results from previous studies suggesting that a balance of excitation and inhibition in cortical networks promotes criticality, thereby maximizing dynamic range and improving input processing^35^ we investigate the effects of varying the E/I ratio, focusing on the dynamic range. It is shown that certain ratios increase the probability of the system operating in the critical regime, thereby extending its dynamic range.

## Results

In the studied network, both excitatory and inhibitory neurons receive a Gaussian input characterized by an identical mean and standard deviation. The input’s mean value effectively maintains the membrane potential below the threshold, yet the non-zero standard deviation or noise can stochastically drive the neurons to the threshold. Once a neuron reaches or exceeds a specific threshold, it fires and then returns to its resting state. This behavior suggests that neurons oscillate between their resting state and activation threshold. In this research, the activation threshold and resting state are represented by numerical values one and zero, respectively. When excitatory neurons fire, they increase the potential of their neighboring connections by an amount *K*, while inhibitory neurons decrease it by the same amount. To investigate balance and criticality, we use two key variables: coupling strength (*K*) and inhibitory percentage (*I*). Specifically, the network displays critical behavior when the coupling strength (*K*) is equal to the critical value (*K*_*c*_). The neurons are distributed on a lattice with periodic boundary conditions and simulations are performed on lattices with *N* = 1600, 2500, and 3600 neurons where *N* = *L×L* for *T* = 2×10^7^ time steps (see Methods section).

Here we first, compare the spike and *PAV* time series generated by the model for different *K* values. The *PAV* is defined as follows:

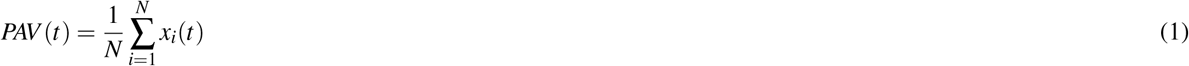

where *N* denotes the total number of neurons within the network, while *x*_*i*_(*t*) refers to the membrane voltage of the *i*^′^*th* neuron at time *t*. Fig. 1 shows spikes and the associated *PAV* for three different control parameters when 20 percent of the neurons in the network are inhibitory (*I* = 20). This illustration shows that the branching pattern becomes apparent when the system operates in the critical and super-critical regimes. In contrast, within the sub-critical regime, all *PAV* fluctuations are suppressed close to the threshold of the neuron. This behavior is a result of the irregularities observed in the spike patterns. The following sections provide detailed explanations on how to determine critical, sub-critical, and super-critical regimes. Additionally, the sections shed light on the contributions of excitatory and inhibitory neurons to the overall network dynamics, with a focus on the order parameter and the analysis of neural avalanches. The first step involves determining the transition type based on varying percentages of inhibition. Next, neural avalanche analysis is conducted to investigate the impact of inhibitory neurons on the observed behaviors.

**Fig. 1.**
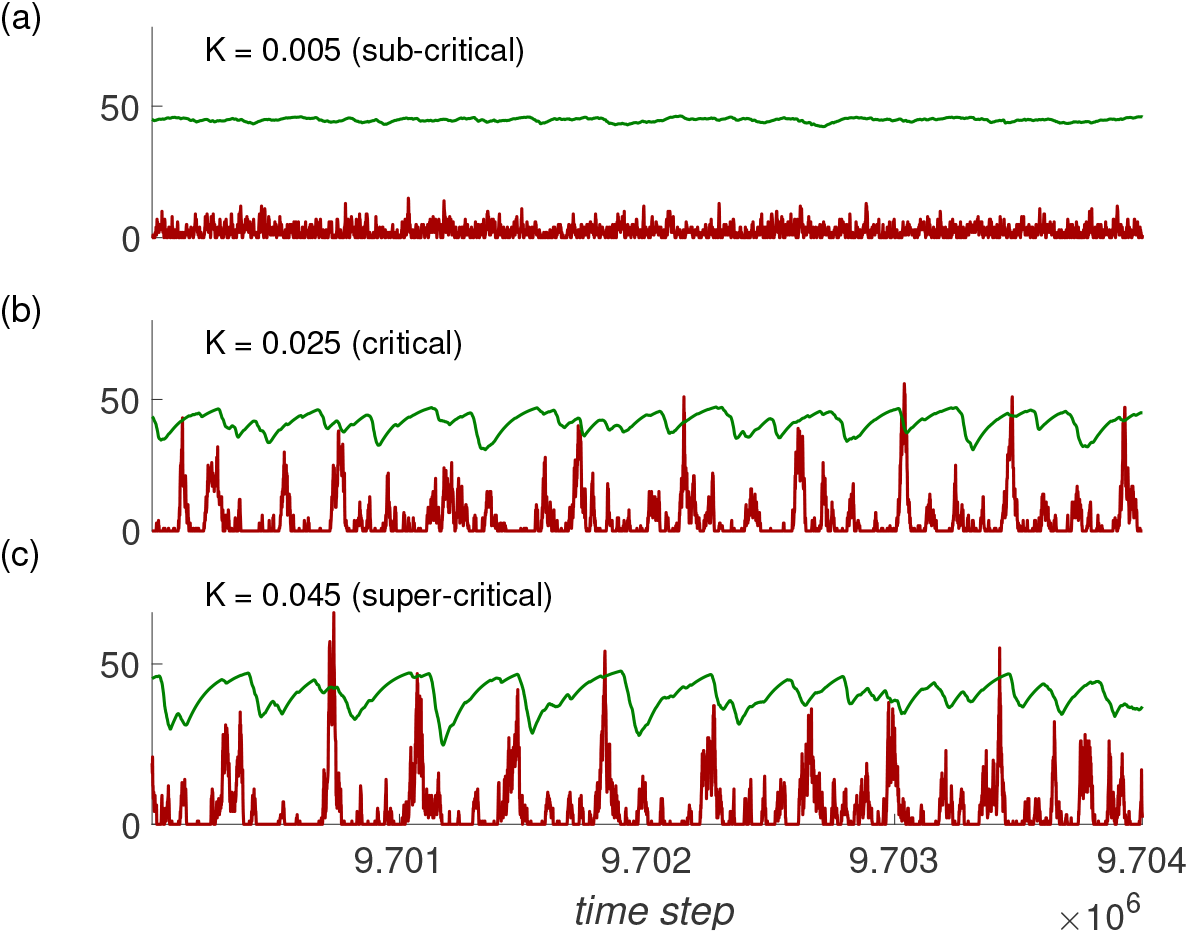
The time series data is presented for spike number (in red) and the product of *PAV* and *L* (*PAV* multiplied by *L*, denoted in green): (a) in the sub-critical state with *K* = *0*.*005*, (b) at the critical point with *K* = *0*.*025*, and (c) within the super-critical domain at *K* = *0*.*045*. The linear’s size (*L*) is 50, which corresponds to a total of 2500 neurons, of which 20 percent are inhibitory.

### Order parameter and dynamical range

This section will begin by reviewing the definition of the order parameter for a pure excitatory neural network^57^. The mentioned definition has a crucial role in quantifying the synchronization within the network. Moreover, by employing the order parameter, we can characterize the critical points where the network transitions between various states of synchronization. The systematic comparison of different states within the network is possible by employing this parameter, particularly when considering various inhibition levels. Such comparative analysis enhances our understanding of how the percentage of inhibitory neurons affects the overall dynamics and helps develop a comprehensive understanding of the network’s behavior. Here, *SPC* measures the connection between the *PAV* and the timing of neuronal spikes. To calculate the *SPC*, three steps will be performed:

1. The instantaneous phase of *PAV* is calculated for each spike in the spike train of every neuron in the network. This is performed using the Hilbert transform, and the results are then averaged to obtain a mean instantaneous phase value for *j*^′^*th* neuron in the network (*M*_*j*_).

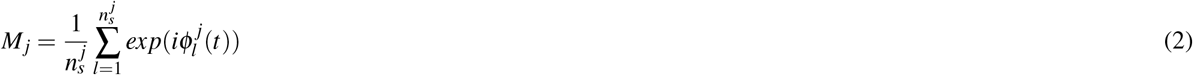

where 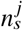 shows the number of spikes for neuron *j* and 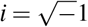. Here, 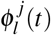 is the instantaneous phase of *PAV* corresponding to the *l*^′^*th* spike of the *j*^′^*th* neuron and is extracted from the Hilbert transform of the *PAV* time series.
2. The phase-locking value (*PLV*) is calculated by averaging over all *M*_*j*_ values^61^.

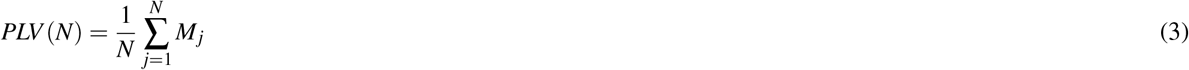

where *N* is the number of neurons in the network (*N* = *L×L*). The *PLV* functions as a synchronization metric. A *PLV* value of zero indicates either a lack of spikes or complete desynchronization (when *n*_*s*_ → ∞), while a value of one (in absolute terms) indicates perfect phase locking and synchronization between the spikes. This step assists in evaluating the alignment of the phases of different spikes, which provides a measure of coherence within the system.
3. The absolute value of *PLV* is used to estimate the *SPC*. This absolute value is used as the order parameter (*m*) to describe the synchronized behavior of the network.

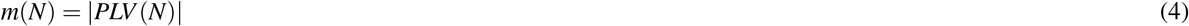

Fig. 2a shows variations of order parameter, *m*, with respect to the control parameter, *K* for a particular lattice size, *L* = 40. It shows that including inhibitory neurons has a dual effect. First, it widens the transitional region between de-synchronized and synchronized states, effectively increasing the range of conditions under which these transitions occur (the horizontal length between the stochastic and synchronized parts of the transition region expands). Second, it leads to a reduction in the value of the parameter *m*, especially for stronger coupling strength at high levels of inhibition. To further investigate, we evaluate this expansion by the dynamic range, providing insight into how the varying ratio of excitatory and inhibitory neurons affects the system’s behavior. While the transition between states remains continuous across all examined ratios, a distinct pattern emerges, specifically at an inhibitory ratio of 20%. To quantify these transitions for different inhibition percentages, we employ the dynamical range, denoted by Δ_*h*_:

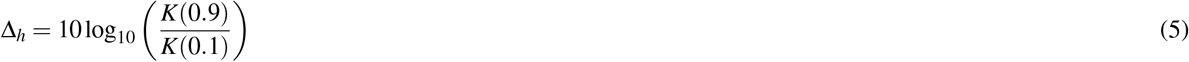

**Fig. 2.**
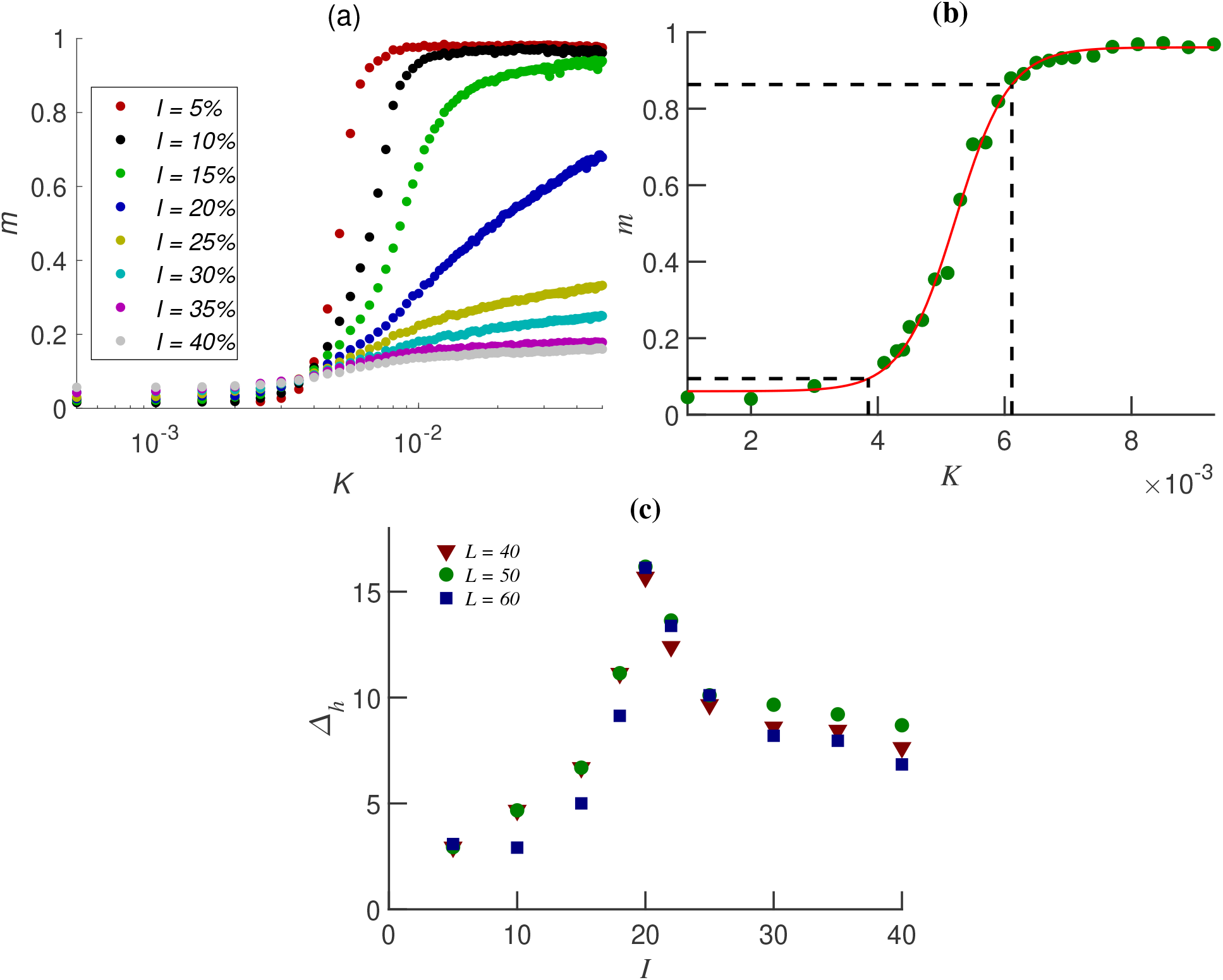
(a) The dependence of the order parameter *m* on the parameter *K* is shown for lattices of linear size *L* = 40 and different inhibition percentages: 5%, 10%, 15%, 20%, 25%, 30%, 35%, and 40%. (b) The plot of *m* as a function of *K*, with vertical lines indicating the values of *K*(0.9) and *K*(0.1) for *L* = 60 and an inhibition percentage of 5%. (c) Analysis of the dynamic range Δ_*h*_ over different lattice sizes and inhibition percentages.

Here, *K*(*x*) is the value of *K* for which the order parameter curve attains a value that is *x* times the maximum of *m*, or more specifically:

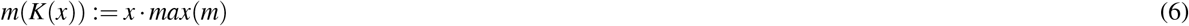

The calculation of the dynamic range for different inhibitory percentages and lattice sizes requires determining the order parameter at higher *K*, where it becomes saturated and independent of *K*, as shown in Fig 2b. The black dashed lines in the figure assist in identifying the key parameters *K*(0.9) and *K*(0.1), which are then used to compute the dynamical range, as given in Equation (5). Additionally, Fig 2c shows the dynamic range as a function of different inhibition ratios. The figure shows that the dynamic range reaches its maximum value for *I* = 20%, a result that is consistent for all lattice sizes examined. The results of the study show that while inhibitory neurons in the system may initially prevent or postpone synchronization, the introduction of an optimal level of inhibition can actually expand the dynamic range of the system. This is particularly noticeable when the control parameter is significantly larger than that of networks with other inhibitory percentages and the system is still not in synchronized phase. Therefore, inhibition performs a subtle function, both restricting and perhaps enhancing the operational flexibility of the system. The depicted transition between different states of synchronization is continuous, highlighting the gradual shift in behavior as the parameters vary. We prove that this system undergoes a continuous transition and critical behavior by using the finite-size scaling theory. This approach provides a better and more detailed comprehension of the system’s behavior. In this context, we examine the interplay between the order parameter, coupling strength, and lattice size, considering the network’s composition of 20 percent inhibitory neurons. Here, the relationship between the order parameter, coupling strength, and lattice size is assumed as follows:

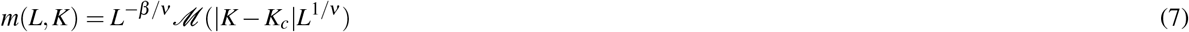

*β* and ν are the critical exponents corresponding to order parameter and correlation length, respectively. *ℳ*(*x*) is the scaling function. For systems of large sizes, the behavior of *ℳ*(*x*) is expected to be as follows:

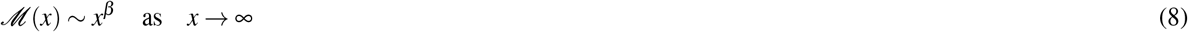

In the infinite size limit when *K* > *K*_*c*_, the following relation is necessary for a continuous phase transition.

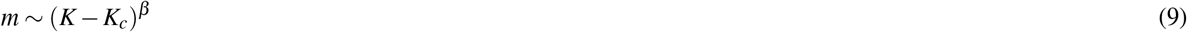

Fig. 3a shows the order parameter as a function of the control parameter parameter *K* for inhibition level of 20% at lattice sizes *L* = 40, 50, and 60. Fig. 3b illustrates the collapsing of the data over a range of lattice sizes for 20% inhibition percentages. The observed behavior shows a distinct class of universality, differing from both mean-field directed percolation with *β* = 1 and ν = 0.5, and directed percolation with *β* = 0.580(4) and ν = 0.729(1) in 2+1 dimention^62^. In more detail, the result suggests that synchronization within the system is suppressed by the inhibitory neurons’ attenuating effect on excitatory activity. The underlying mechanisms of this effect can be understood in terms of the synchronized activity between excitation and inhibition. Here, inhibitory inputs can balance the excitatory input received by an excitatory neuron when both types of neurons are active simultaneously. As a result, while synchronization between excitatory and inhibitory neurons may initially prevent the synchronized phase within the system, which is measured by the m-value, the implementation of an optimal level of inhibition can indeed expand the dynamic range. This effect is especially noticeable when the control parameter is significantly large and the system has not yet been synchronized. Therefore, it might be a result of a different phase between excitatory and inhibitory activity.

**Fig. 3.**
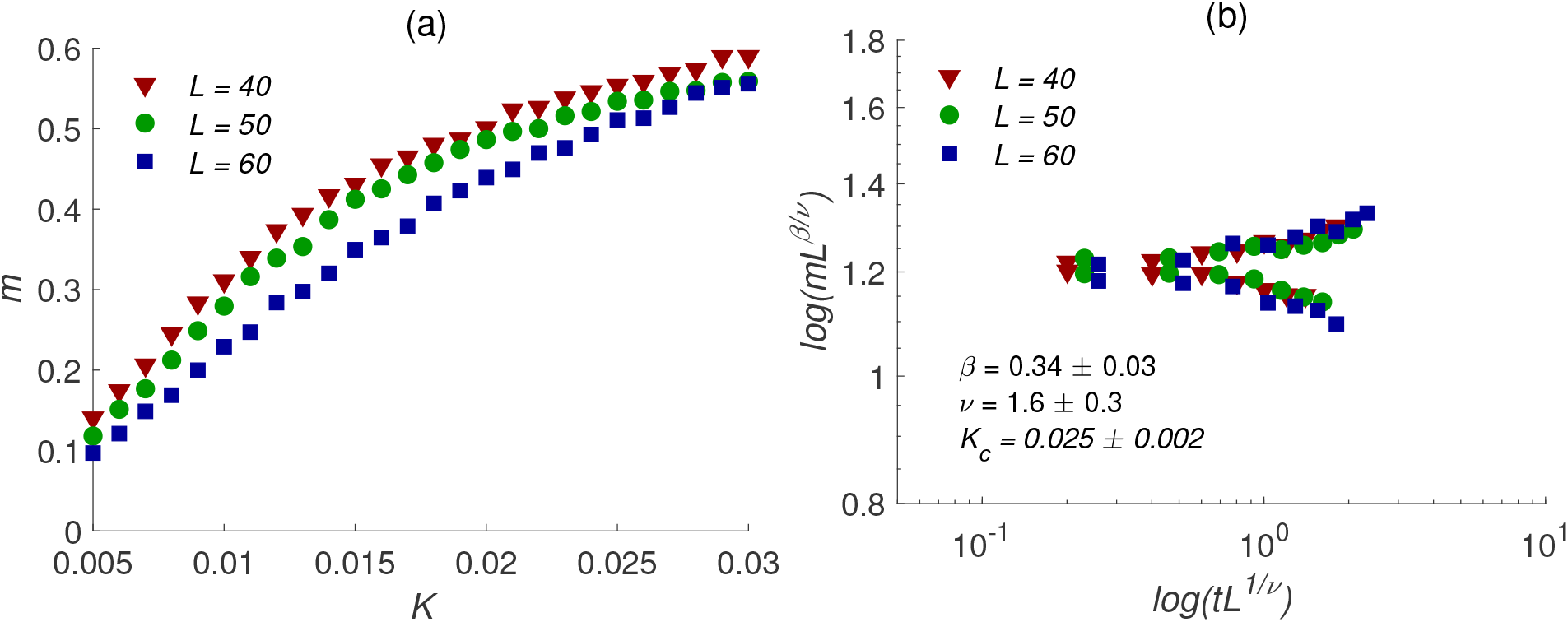
(a) Variation of the magnitude of the order parameter (*m*) with parameter *K*, plotted for lattices of linear sizes *L* = 40, 50, 60 and inhibition ratio 20% around the critical point. (b) Rescaled *m* − *K* curves showing collapse for different lattice sizes and inhibition ratio 20%. Note: 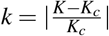

To evaluate the effectiveness of inhibitory neurons across the dynamic range, it is imperative to study E/I ratio in different network topologies. To compute the dynamic range, this study employs a small-world network model with variable rewiring probabilities^63^. This is consistent with the concept that the brain probably functions as a small-world network, although further research and discussion continue to support this claim^64–66^. Figure 4a shows the dependency of the dynamic range over the range of rewiring probabilities. As it is shown in Figure 4b, when *p*_*r*_ = 0, the dynamic range will be maximized when there is 20 percent inhibition. This shows the results can be universal, and the choice of small-world network has the benefit of a wider dynamic range in comparison to lattice. However, this raises the question of whether the network evolution objective function is the expansion of the dynamical range or whether this is a subresult of a more general objective function.

**Fig. 4.**
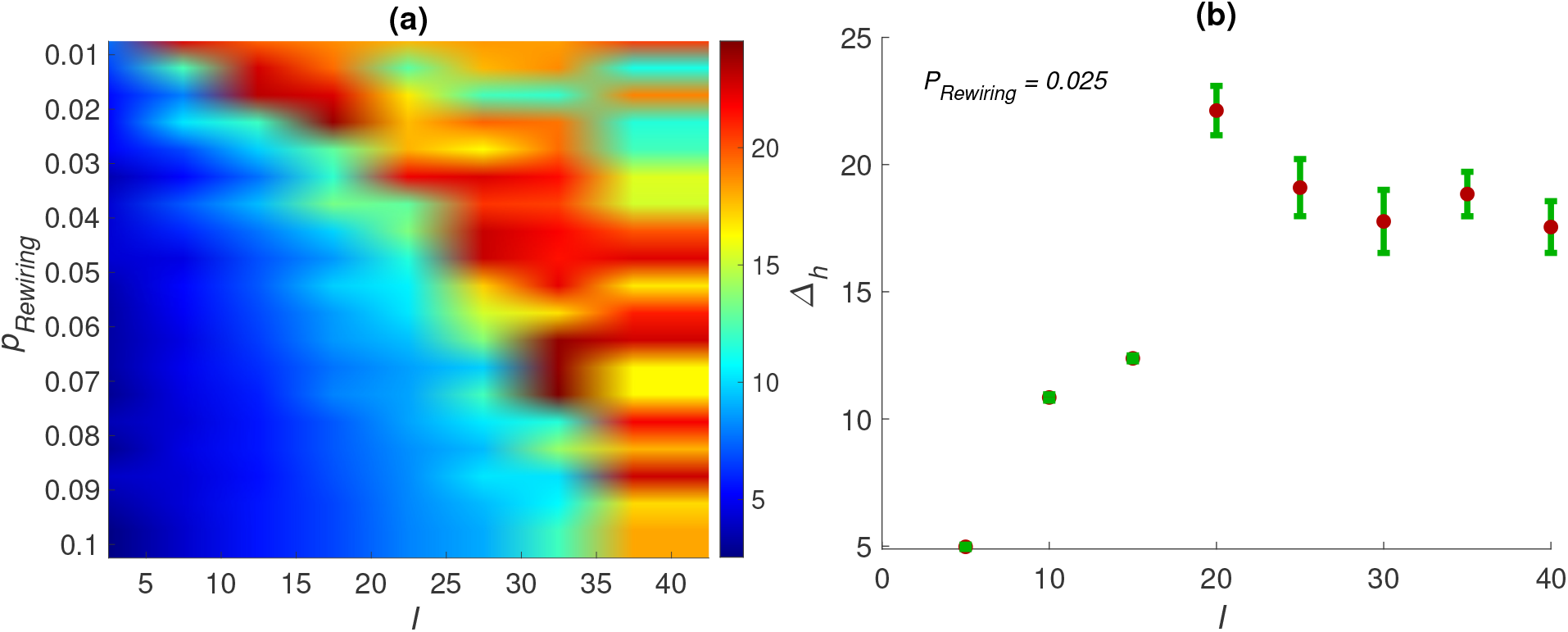
(a) Variation of the dynamical range under different inhibition percentages and a spectrum of rewiring probabilities. (b) Analysis of the dynamical range under different inhibition percentages with a constant rewiring probability of 0.025. Initially, these models use a Watts-Strogatz network framework defined by a 4-degree complete graph with 2500 neurons.

### Inhibition effects on neuronal avalanches

In the current study, the introduction of inhibition into the network resulted in the same type of transition from desynchronized to synchronized behavior in a pure LIFM^57^. However, this led us to question how inhibition operates within the coupling domains, leading to the emergence of criticality, a region we specifically call the critical domain. Our recent studies investigated the emergence of criticality using temporal complexity and avalanche analysis^67–69^ in a leaky integrate-and-fire model following finite-size scaling^57^.

In this study, we perform neural avalanche analysis to determine the possibility of criticality emergence by considering different percentages of inhibition in the system. In more detail, an avalanche is characterized by two main features: its size, represented by *S*, and its lifetime, commonly known as duration, represented by *T* (see the Methods section). Neural systems exhibit diverse sizes and durations, where their probability distributions follow a power-law pattern^32^. This behavior is characterized by a single universal scaling exponent: *τ* for size, represented by *p*(*S*) = *S*^−*τ*^, and *α* for duration, represented by *p*(*T*) = *T* ^−*α*^ which follwos a shape collapse in cultured slices of cortical tissue^38^.

Despite the power-law behavior of avalanche size and duration, demonstrating criticality behavior in avalanche data requires universal scaling^70^. Therefore, we perform a scaling analysis over different network sizes and different inhibition percentages. According to the single scaling theory, in the case of a network with a size of *L*, the probabilities are expressed as follows^70^:

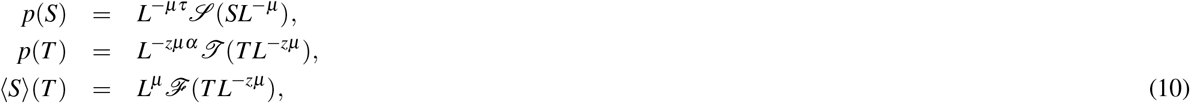

where *µ* is the scaling exponent of the size of avalanches versus the linear size of the lattice and *z* is the dynamical exponent which determines the scaling relations of the duration and size of the avalanches. Here ⟨*S*⟩ (*T*) is the average of avalanche size conditioned on a given duration^70^. Here *ℐ*(*x*), *ℒ*(*x*), and *ℱ*(*x*) represent universal functions. Their behavior in the large *x* limit (*x* → ∞, corresponding to small *L*) is described as follows:

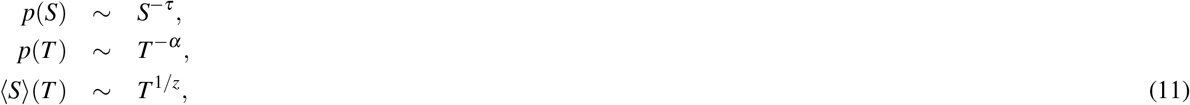

Moreover, the scaling theory requires the following exponentiation relationship^38,71^:

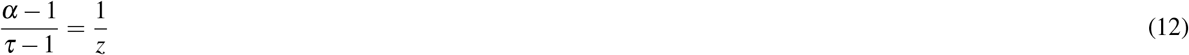

in which the mean field prediction for the scaling exponents are *τ* = 3*/*2, *α* = 2.0, and 1*/ 𝓏* = 2.0^70^. We perform this analysis on the neural avalanche patterns for networks in which 5%, 10%, 15%, and 20% of the neurons are inhibitory.

Fig. 5 shows neural avalanches for inhibition percentage of 20 that follow a scaling relationship and collapse into a single function. A value of ≈ 2.0 for *µ* suggests that the avalanche size grows directly to the lattice size.

**Fig. 5.**
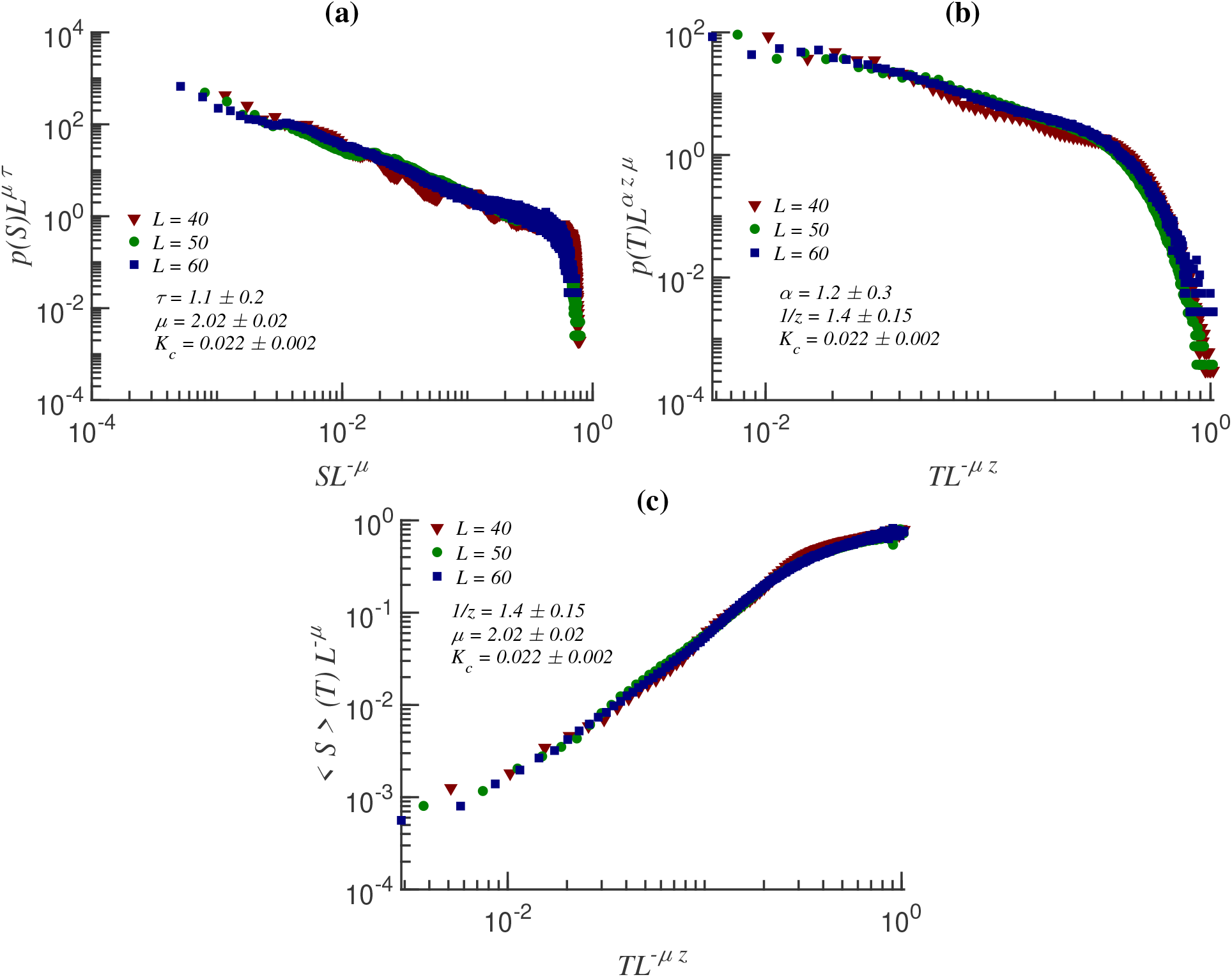
The figure displays the collapse of avalanche data through the scaling relation. Panel (a) display the plotted avalanche sizes whereas panels (b) present the duration probabilities *p*(*T*) and the conditional average of avalanche size versus duration shown as (c). The results are presented at the critical point *K* of square lattices with linear sizes of *L* = 40, 50, and 60, and inhibitory percentages of *I* = 20%. The slope of maximum avalanche size vs. lattice size was used to obtain *µ*.

Moreover simulations show that inhibitory neurons broaden control parameter’s domain, corresponding to critical behavior (Table. 1). Neural avalanches show power-law behavior over a broad range of *K* values, especially when the inhibition percentage *I* is set to 20%. Furthermore, the parameters that describe the phase transitions vary for each inhibitory percentage.

**Table 1.** Exponents for data collapsing for different inhibitory percentages.

| | $I = 5$ | $I = 10$ | $I = 15$ | $I = 20$ |
| --- | --- | --- | --- | --- |
| $\nu$ | $1.05 \pm 0.05$ | $1.85 \pm 0.15$ | $1.78 \pm 0.05$ | $1.6 \pm 0.3$ |
| $\beta$ | $0.32 \pm 0.17$ | $0.55 \pm 0.05$ | $0.33 \pm 0.03$ | $0.34 \pm 0.03$ |
| $K_c$ | $0.0052 \pm 0.0001$ | $0.0072 \pm 0.0005$ | $0.0115 \pm 0.001$ | $0.025 \pm 0.002$ |

## Discussion

Inhibitory neurons and neuronal avalanches play a crucial role in understanding complex neural networks. In our previous research, we studied a purely excitatory network and showed a dynamic transition of neuronal behavior from an asynchronous state to a fully synchronized state, with a distinct and complex critical phase between these two phases. Furthermore, we quantified this transition using the Kuramoto order parameter.^57^

Moreover, previous research emphasizes the importance of the E/I ratio in network dynamics, especially at high connectivity levels^72^ and have shown that the local excitation-inhibition ratio strongly influences whole-brain dynamics, the structure of spontaneous activity and information transmission^73^.

In this context, our primary focus in this study was to examine the impact of different E/I ratios on the emergence of criticality and phase transition. It is shown that an optimal ratio (20%) facilitates critical behavior in the network. Inclusion of 20% inhibitory neurons expands the coupling parameter domain in which neural avalanches satisfy the scaling relation, increasing the probability of finding the network in a critical state and maximizing the dynamic range.

The observed effects might be associated with a subset of inhibitory neurons that are in the same phase as excitatory neurons. This prevents synchronization at larger connectivity values (*K*) compared to networks with different inhibitory neuron percentages. As a result, the difference between *K* that leads to synchronized and desynchronized states expands, resulting in a more extensive range of dynamic behavior. However, excessive levels of inhibition can prevent the emergence of synchronization. In such scenarios, the system can be pushed into a permanent desynchronized state. This might emphasize the intricate balance necessary for optimal control parameter between excitatory and inhibitory neurons.

The study results, although they concentrate on a particular computational model, also provide insight into broader principles that may apply to other network topologies, especially in the context of criticality, which manifests itself as transitions between desynchronized and synchronized states. Fundamentally, different levels of inhibition can significantly modify the network’s dynamic range. The degree of inhibition could, for example, limit the dynamic range at low percentages of inhibition and possibly lead to a lack of transitions in networks dominated by inhibitory activity. It is possible that for another topology, another optimal inhibitory percentage exists, maximizing the dynamic range. This indicates that modulation of neural activity via E/I ratios is a complex process, with different network topologies leading to different outcomes. However, the evolutionarily precise objective function that results in this specific E/I ratio in the cortex is still a matter of further investigation. However, this objective function might drive neural systems toward critical points^74,75^, the dynamic behavior of the order parameter despite synaptic plasticity needs to be investigated.

Taken together, although the results of this study are derived from a model with a defined set of parameters and lattice structure, the identified patterns in the model suggest a likely framework for understanding how E/I ratios might modulate cortical scale-free dynamics and contribute to the broader discussion of brain dynamics.

## Methods

### Model description

In this study, we employ the Leaky Integrate and Fire Model (LIFM) ^57,67–69,76,77^. The dynamic of the membrane potential of a single neuron *x*_*i*_(*t*), which is residing on a two-dimensional square lattice of linear size L with the periodic boundary condition, is

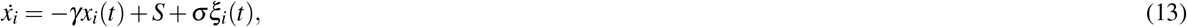

where 1*/γ* is the membrane time constant, assumed to be the same for both excitatory and inhibitory neurons for simplicity. *S* denotes a constant that drives the membrane potential to the steady-state *x*_*i*_ = *S/γ* in the absence of the third term on the right side of the equation, and *ξ*(*t*) is a continuous Gaussian noise having a mean of zero and a variance of one, as defined by

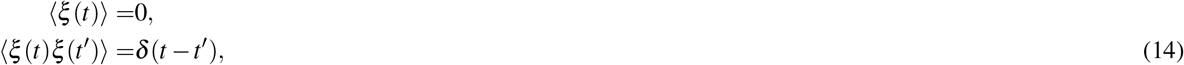

where σ represents noise intensity, which can drive *x*(*t*) to the threshold, in this case *S < γ*, and leads the neuron to fire. In this context, (*S* + *σξ*_*i*_(*t*))*/γ* can be interpreted as the input to each neuron. We randomly distribute *N*_*E*_ excitatory and *N*_*I*_ inhibitory neurons, *N* = *N*_*E*_ + *N*_*I*_, on the lattice. In principle, inhibitory neurons could have two types of inhibitory-inhibitory, inhibitory-excitatory, and excitatory-inhibitory connections. Here, however, there is no link between inhibitory neurons, i.e., no inhibitory-inhibitory connections exist.

The initial membrane potential for all neurons in the network is a random value between 0 and 1 from a uniform distribution.

Considering the reset mechanism by setting the threshold to one; whenever the membrane potential reaches or passes the threshold, i.e., *x* = 1, it jumps back to the resting state, *x* = 0. If the firing neuron is inhibitory, *K* (control parameter) is subtracted from the membrane potential of its adjacent neurons, and if it is excitatory, *K* is added to the membrane potential of its adjacent neurons. The quantity *K* > 0 controls whether the system is in a sub-critical, critical, or super-critical state. We refer to *K* as a control parameter in which *K*_*c*_ leads to power-law behavior in the model. The simulations in the current paper are based on the following parameters: *S* = 0.001, *γ* = 0.001005, and σ = 0.0001, and the lattice with linear sizes of *L* = 40, 50, and 60. The integration time step for the numerical solution was Δ*t* = 0.01 (Euler method), and the simulations lasted for 2 × 10^7^ steps.

Implementing the above parameter concludes that only noise and control parameter make neurons pass or reach the threshold. However, for the case of a single neuron, as illustrated in Fig. 6, the neuron preferentially fires within a particular time called *T*_*p*_, the peak in the figure, and spike time intervals follow the Poissonian distribution. Considering a single neuron in the network leads to a change in the amount of this peak per phase with different *K*_*c*_.

**Fig. 6.**
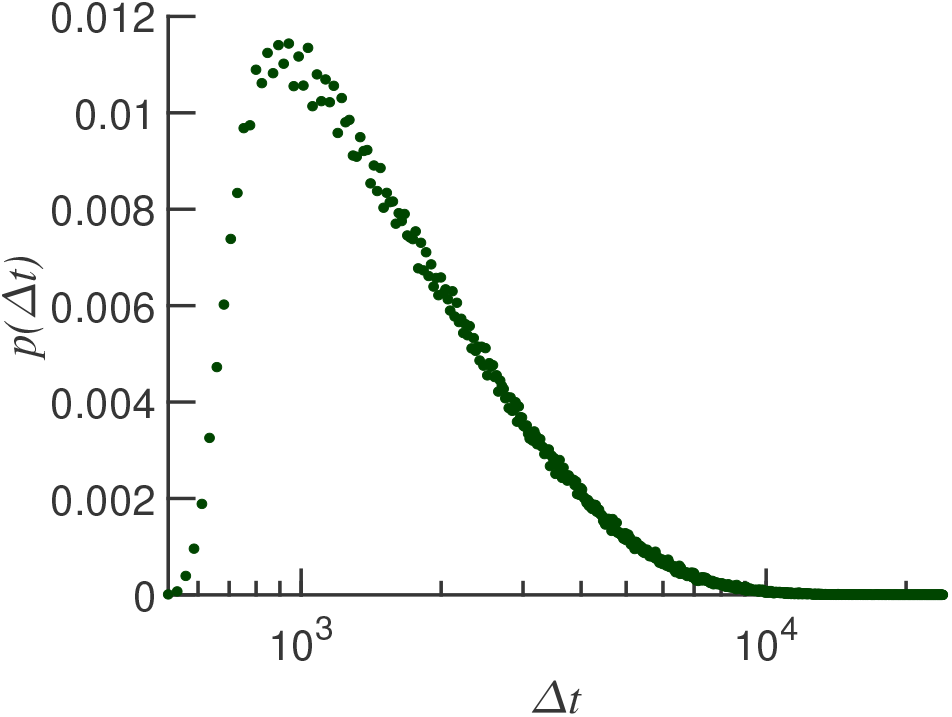
The probability distribution function for the spike time intervals of a single neuron. The simulation time is 10^9^ steps.

### Neural Avalanches

The term “neural avalanche” refers to the sequential propagation of neuronal firing activity within a network in a correlated manner, initiated by firing a single neuron or a small group of neurons. This activity can cause a network to behave in a synchronized manner. To identify neuronal avalanches, we identify time points where there is an absence of firing activity across all neurons in the system for a continuous duration of at least five time steps (*δt* = 5). As shown in Fig. 7, the initiation of an avalanche is marked by the first firing event within the network, indicated by the dashed lines. In this context, a neural avalanche is defined by two key parameters: its size, represented by the number of spikes within the avalanche, and its duration, representing the time span from the initial firing to the last firing event. To comprehensively analyze these neural avalanches, we assess the probability of their occurrence over different sizes, *P*(*S*), and durations, *P*(*T*).

**Fig. 7.**
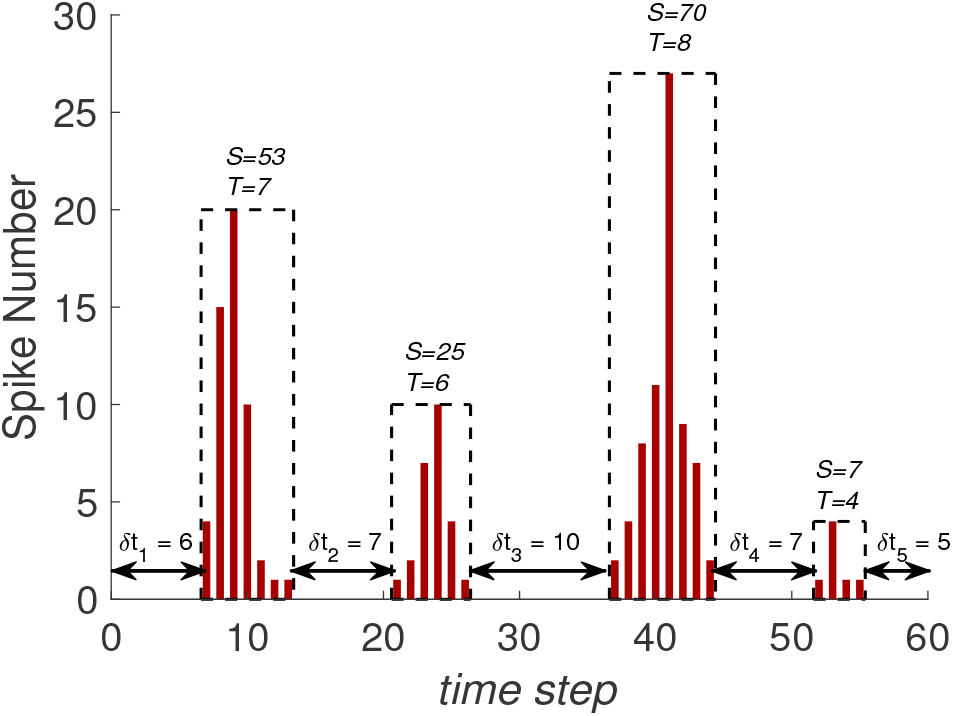
A schematic representation of the process for calculating neuronal avalanches. The steps include: 1. Simulating Neuronal Activity: Neuronal firing patterns are generated and observed over time within the model. 2. Identifying Silent Periods: Silent periods where there is no firing activity across all neurons for at least five time steps (*δt* = 5) are identified. 3. Detecting and Analyzing Avalanches: The regions between consecutive silent periods are considered avalanches. The size of an avalanche is determined by the number of spikes during its duration. The figure demonstrates how the sequential propagation of neuronal firing activity within a network results in the formation of distinct avalanches of various sizes and durations.

## Data Availability

The simulation codes used in this study are available upon request. Please contact the corresponding author for access.

## Acknowledgements

All authors gratefully acknowledge financial support from their affiliated institutions.

